# Inhibition of Herpes simplex virus and Vesicular stomatitis virus Proliferation by *Lactiplantibacillus planterum* and star anise Extract Is Associated with Induction of *MX* gene expression

**DOI:** 10.1101/2023.01.11.523569

**Authors:** Dalia Elebeedy, Asmaa Sayed Abdelgeliel, Aml Ghanem, Ingy Badawy, Bayan H. Sajer, Alya Redhwan, Mashail A Alghamdi, Mohamed R. Eletmany, Marwa El-Sayed

## Abstract

**Purpose:** There is an immense need to develop new antiviral compounds that are effective and have little side effects. Natural compounds provide a good candidate that is gaining popularity. Probiotics as *Lactiplantibacillus plantarum* and plant-derived medicines as star anise extract have been investigated to treat viral diseases, cancer, and inflammation. This work aimed to investigate the antiviral effect of probiotics (*Lactiplantibacillus plantarum*) and (star anise) against Herpes simplex virus type 1(HSV-1) and Vesicular stomatitis virus (VSV).

**Patients and methods:** *Lactiplantibacillus plantarum* and star anise extract had been prepared and tested against vero cell lines for cytotoxicity assay. The antiviral effect of *L. plantarum* and star anise against HSV-1 and VSV was evaluated by titration calculation (TCID50). The expression of *MX* genes were measured in infected cells before and after treatment with *L. plantarum* and star anise.

**Results:** We found that *L. planterum* is more effective against *HSV-1* and caused 2.5 log reduction in virus titre while star anise extract is more effective against VSV and caused 1.25 log reduction in virus titre. Evaluation of *MX* genes expression revealed higher expression in HSV-1 infected cell treated with *L. planterum* and VSV infected cells treated with star anise extract.

**Conclusion:** *L. planterum* and star anise could be useful antiviral natural compounds that have minimal side effects

**Plain language Summary:** Both Herpes simplex virus type 1(HSV-1) and Vesicular stomatitis virus (VSV) caused infection characterized by lytic lesions. *Lactiplantibacillus plantarum* and star anise had shown promising antiviral effects. So, we investigated the antiproliferative effect of *L. plantarum* and star anise against HSV-1 and VSV by assessment of virus titre before and after treatment and assessment of *MX* gene coding for antiviral proteins. We found that *L. plantarum* and star anise could inhibit proliferation of HSV-1 and VSV and increase expression of *MX* gene. *L. plantarum* and star anise could be good antiviral candidate.

## Introduction

HSVs are DNA viruses causing infections in various host species and characterised by lytic lesions.^1^ HSVs has a tropism for epithelial cells and neurons of the peripheral nervous system (PNS) and humans are the only natural hosts.^2^

Herpes simplex causes painful, burning, or pruritic clusters of vesicles on the lips, oral mucous membranes, genital region, or other areas of the body.^3^

Herpes simplex virus type1 (HSV1) is virus that usually causes oral herpes such as cold sores and fever blisters and it can cause some forms of genital herpes [4]. About 60%–80% of people throughout the world are infected by HSV1.^5^

Vesicular Stomatitis virus is RNA virus that affects livestock forming blister-like lesions. Human infections cause febrile illness and are associated with exposure to infected animals.^6^

Probiotics are beneficial microorganisms found in food and supplements.^7^ *Lactiplantibacilli* are Gram-positive rods that breakdown polysaccharides anaerobically and produce lactic acid.^8^ *Lactiplantibacillus plantarum* exhibits an antiviral effect against HSV-2.^9^

Star anise is an Asian cooking ingredient. It has an antiviral effect against some viruses such as HSVs. Star anise oil, rich in transanethole, revealed a high selectivity index of 160 against HSV.^10^

Our study aims to investigate the antiviral effect of *Lactiplantibacillus plantarum* and star anise against HSVs and VSV.

## Material and methods

### Preparation of probiotic

*Lactiplantibacillus planterum* was obtained from ATCC (14917). Cells were grown in MRS (Man Rogosa Sharp) medium at 37 °C, for 24 h in shaking incubator. When the media became turbid, it was centrifugated to precipitate the cells and the supernatant was taken. The pellet was taken and saline was added to it and then centrifuge it to remove any residuals. The pellet with overnight bacterial culture was sonicated after adding saline using the intermediate sonication horn of a Sonic 300 Dismembrator Fisher Scientific Co., NJ, and USA at 60% intensity to release the intracellular compounds. The horn is 9.5 mm in diameter, was placed in a 15-ml polypropylene tube containing 5-6 ml of cell suspension in PBS. Cooling was achieved by placing the polypropylene tube in an ice water bath while sonicate the samples. After centrifugation the supernatant was sterilized by syringe filter used.

### Determination of total protein in Lactiplantibacillus planterum supernatant and cell sonicate

Blank tube containing 0.2 ml of 0.85% Sodium Chloride Solution was prepared. 0.2 ml of a test sample solution was prepared and to each tube 2.2 ml of the Biuret Reagent was added and left at room temperature for 10 minutes. 0.1 ml of the Folin and Ciocalteau’s Phenol Reagent were added to each tube, mixed each tube well immediately after addition and left at room temperature for 30 minutes. Solutions were transferred to microplate and absorbance was measured within 30 minutes. The protein concentration (mg/ml) was determined from the standard curve.

### Extraction of star anise

The star anise was purchased from local market and then cleaned. The samples were chopped and finely ground in a blender. The ground samples were then extracted with solvents either in cold water at 30°c for 4 h or in hot water at 100°c for 15 min. After the extraction was terminated the samples were filtered through linen filter cloth. All filtrates were concentrated in a rotary evaporator. Samples were kept in capped dark glass bottles at –20°c till use. Voucher specimens were deposited at the herbarium of South Valley University, Egypt (unregistered herbarium) with S/N number.

### Determination of total carbohydrate

Working standard solutions of glucose was taken in boiling tubes. 1ml of 5% Phenol and 5ml of 96% Sulphuric acid was added in each tube and shook well then, all the tubes were placed in water bath at 25-30°C for 15 minutes. Optical density of each tube was taken at 490nm with spectrophotometer. The whole process was repeated with 0.2ml of different samples and the optical densities were measured.

### Cell line preparation

The Vero cell line was supplied by from VACSERA. Vero cells were cultured in MEM medium (Minimal Essential media) with 5% Fetal Calf Serum (FCS) and incubated at 37°C in a 5% CO2. After 24hr. incubation the media was discarded and then trypsin was added to the cells to separate the attached cells and then the trypsinized cell was incubated for 30mins to make complete dissociation. After that the serum added to the flask to inhibit trypsin activity. 10ml of medium was added to the cells which were counted and distributed in 96 well ELISA plate. The plate was incubated for 24hr at 37 °C in a 5% CO2 atmosphere.

### Cytotoxicity assay for star anise extract, Lactiplantibacillus planterum supernatant and cell sonicate

The samples were divided into 4 tube, each contained 300μL from star anise extract in the first tube, probiotic *L. planterum* (cell sonicate) in the second, probiotic *L. planterum* (cell supernatant) in the third, combination of the star anise extract (100μL), *L. planterum* (cell supernatant (100μL) and cell sonicate(100μL) in the fourth. Cytotoxicity assay was performed in 96 well plate. 200μL was added from star anise and/ or probiotic from four tube samples. Two-fold dilution was done then 100μL from the Vero cell line suspension was added in every well. The plate was incubated for 24hr at 37°c. After 24hr the media was discarded and the monolayer with adherent was washed phosphate buffered saline (PBS, pH 7.4). Cell viability was evaluated by the MTT colorimetric technique. In each well 200 μl of MTT was added to each well then, the plates were incubated for 6-7hr. 100 μl of isopropanol or DMSO was added to the wells to solubilize MTT crystals and then the optical density of each well was determined by the 595 nm using a Micro plate reader (Biotek, USA). Each experiment was done in triplicate.

### Viral assay

96 well tissue culture plate was prepared by adding 100μL from Vero cell line and 100μL from medium (MEM medium), the plate was incubated for 24h in 37°c. The media was discarded and about 100μL from the safe concentration of star anis and *L. plantarum* for was added in the plates and incubated for 24 hr. Three 96 well tissue culture sterile plate was prepared; one of them for (HSV-1), the second for (VSV), the third for the control. Ten-fold serial virus dilutions were prepared then 100μL of dilutions were added to 96 well plate and incubated for one and half hour at 37 °C. After adsorption, 200 μl/well MEM medium with 2 % FBS were added. All plates were incubated at 37 °C and 5 % CO2 then, observed daily for cytopathic effect (CPE). Endpoints estimation was made after 48 h. Virus Titration calculation (TCID50) was calculated use Reed and Munch method for calculation then the difference between virus’s initial concentration and concentration after treatment was calculated. The titration was repeated for three times.

### Analysis of MX gene expression

Max-Discovery RNA Isolation Kit was used to extract equal amount of total RNA from cells of three independent experiment. RNA concentration was determined using a NanoDrop 2000 spectrophotometer (Thermo Fisher Scientific), and 2 μg of RNA was used to generate cDNA using the RT2 First Strand Kit (#330401, SABiosciences, a Qiagen Company). In a fluorescent temperature cycler, each RT reaction solution was amplified in 25 μl of SYBR Premix Ex Taq II (Takara Bio, Inc.) containing 0.2 μM of each primer. The fold change of expression relative to untreated control cells was calculated using the data analysis program.

### Statistical analysis

Data were statistically assessed utilizing IBM Statistical Package for the Social Sciences (SPSS) version 25. Laboratory characteristics were recorded as means and standard deviation for continuous data. Post hoc Bonferroni correction test showed similarities among all groups. *P* value was considered significant if < 0.05.

## Results

### Total proteins of *L. planterum* and total carbohydrate of star anise

The concentration of total protein in probiotic was measured as the proteins are the main component of probiotic which has effect on virus replication (Figure 1). The total protein in *L. planterum* supernatant was the highest (26 mg\mL) and lowest in the cell sonicate (11 mg\mL). The concentration of total carbohydrate in star anise was measured as the carbohydrate is the main component of star anise which has an effect on virus replication. The total carbohydrate in 10% concentration of star anise extracted with water was 5.64 mg\mL.

**Figure 1.**
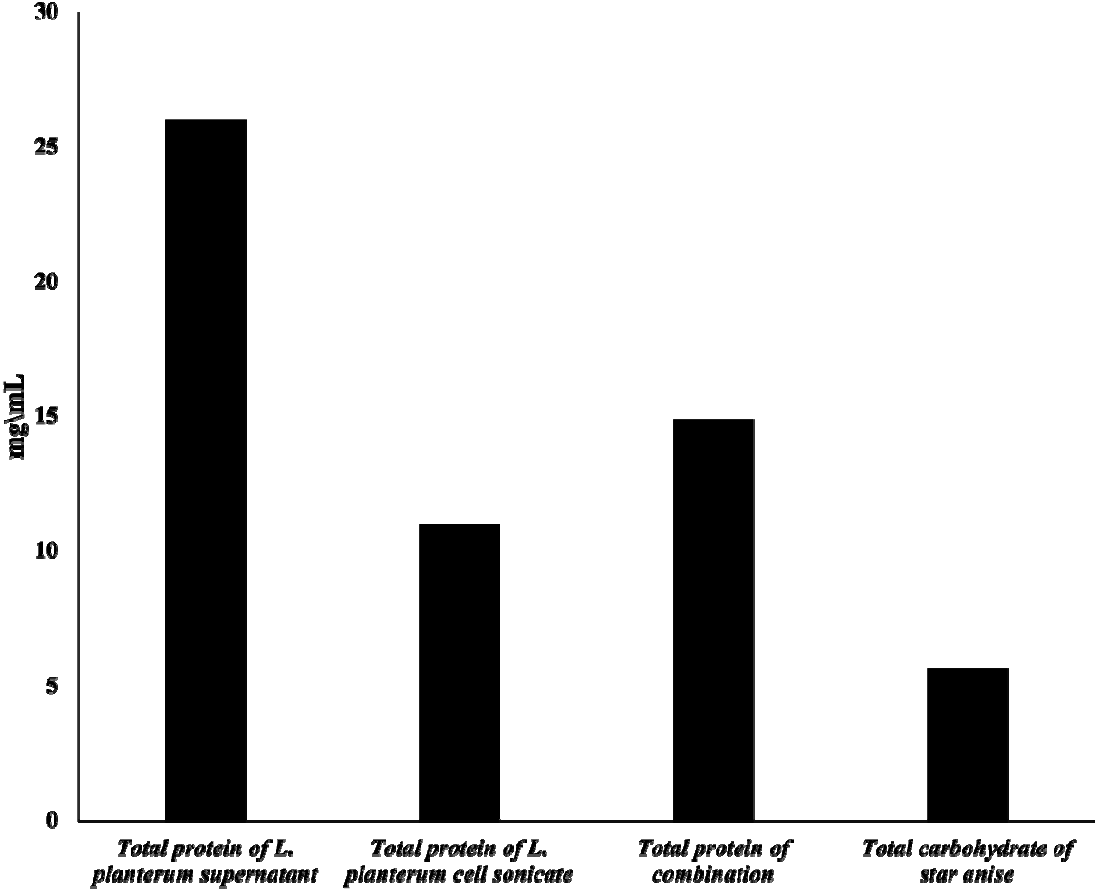
The total protein in probiotic and total carbohydrate in star anise.

### Cytotoxic effect of *L. plantarum* and star anise on Vero cell lines

The cytotoxicity of *L. plantarum* and star anise was measured on vero cell lines (by calculating viability percentage to determine the working safe concentration. Figure 2 represented the IC50 of vero cell line after being treated with *L. plantarum* and star anise. The working safe concentration for star anise was 2.8 mg/mL, 5.5 mg/mL for *L. planterum* cell sonicate, 6.50 mg/mL for *L. planterum* supernatant, 1.86 mg/mL for the combination.

**Figure 2.**
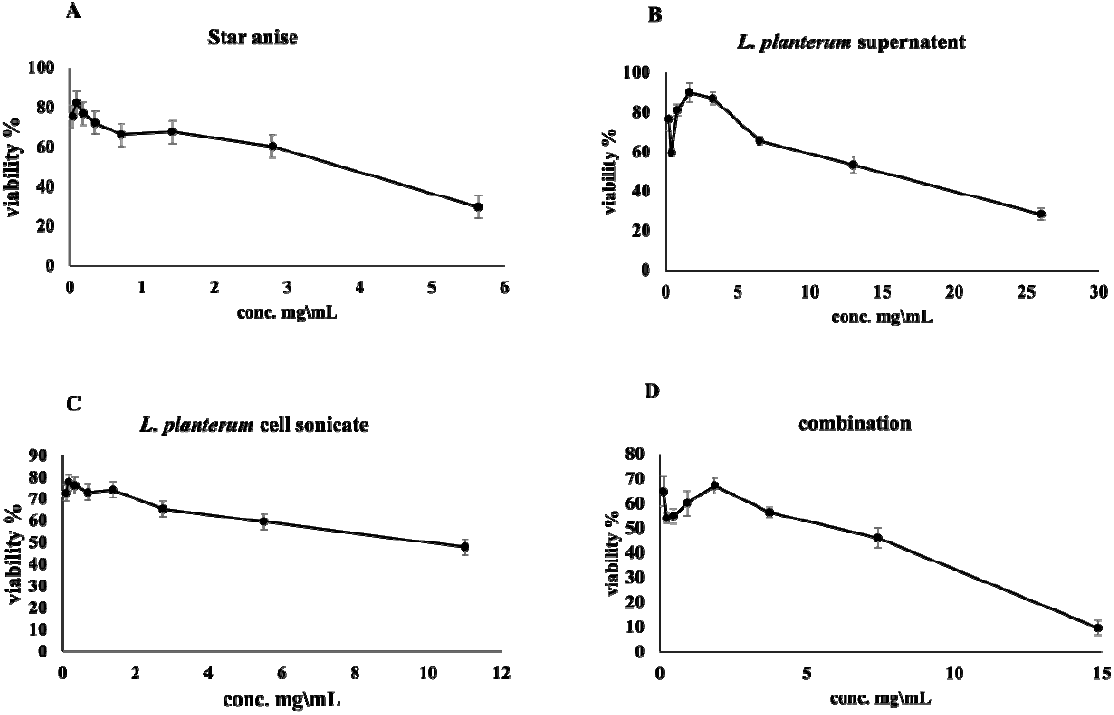
The cytotoxicity test was used to determine the safe concentration of *L. planterum* and star anise on vero cells. A: viability after treatment with star anise. B: viability after treatment with *L. planterum* supernatant. C: viability after treatment with *L. planterum* cell sonicate. D: viability after treatment with combination. All data represent mean ± SEM. Results are the mean of three independent experiments.

### Assessment of of *L. plantarum* and star anise effect on viral replication

We measured reduction in virus titre after treatment with *L. plantarum* and star anise. Log reduction of HSV-1 by star anise or combination is 1.25 while *L. planterum* cell sonicate or *L. planterum* supernatant caused 2.5 log reduction. Log reduction of VSV by star anise extract is 1.25, 0.25 for *L. planterum* cell sonicate, 0.75 *for L. planterum* supernatant and the combination (Figure 3).

**Figure 3.**
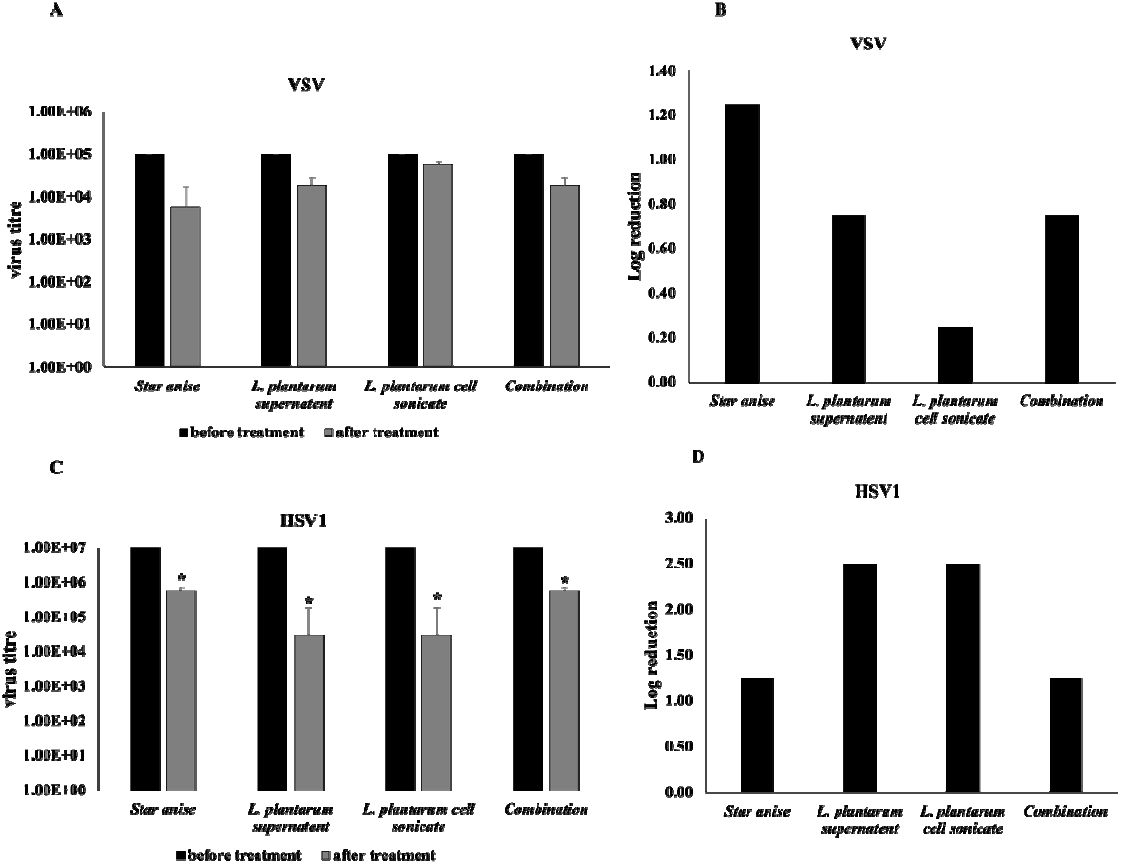
Virus assay before and after treatment with *L. plantarum* and star anise. A: virus titre of VSV before and after treatment. B: Log reduction of VSV titre after treatment. C: virus titre of HSV-1 before and after treatment. D: Log reduction of HSV-1 titre after treatment. All data represent mean ± SEM. Results are the mean of three independent experiments: *p < 0.05 versus before treatment.

### Assessment of *MX* genes expression in Vero cell line after treatment with *L. plantarum* and star anise

We measured the level of *MX* genes expression in Vero cell after treatment with *L. plantarum* and star anise. Figure 4 showed the number of CT of *MX* gene expression after treatment relative to untreated control of Vero cell line in case of HSV virus. Vero cells infected with HSV and treated with *L. planterum* cell sonicate had the highest level of *MX* gene expression. Vero cells infected with VSV and treated with treated with star anise had the highest level of *MX* gene expression.

**Figure 4.**
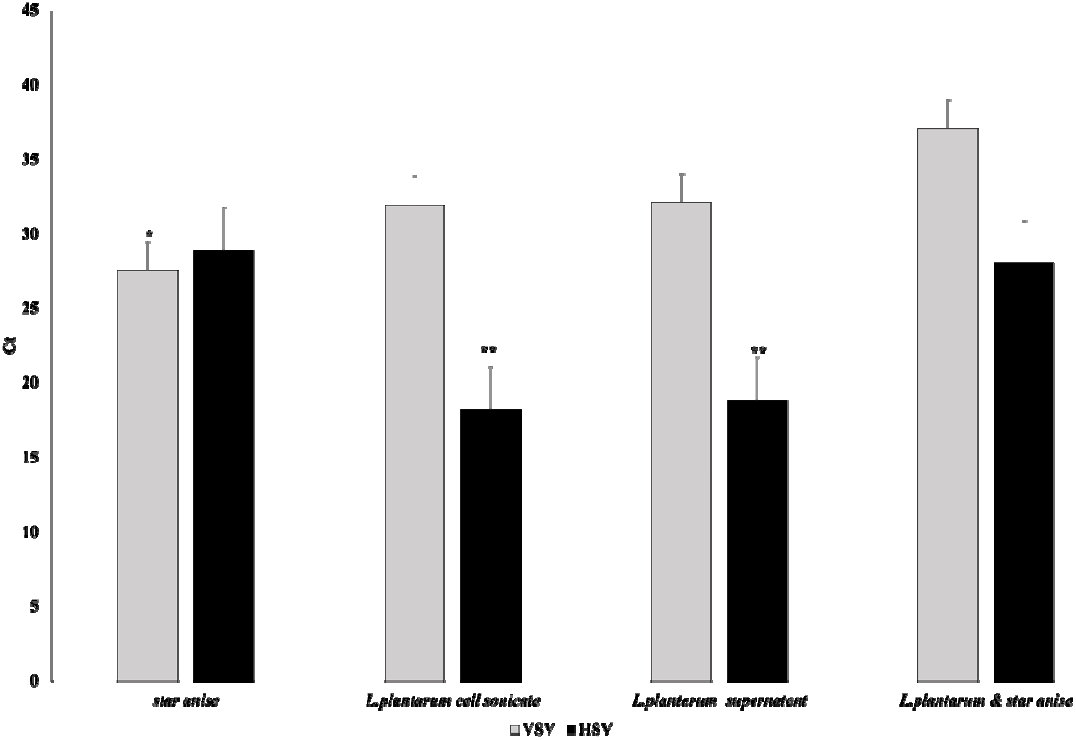
Assessment of *MX* genes expression in HSV and VSV infected vero cells after treatment with *L. plantarum* and star anise. All data represent mean ± SEM. Results are the mean of three independent experiments: *p < 0.05 versus *L. plantarum* and the mixture. **p < 0.05 versus star anise and the mixture

## Discussion

*Lactiplantibacillus plantarum* has versatile therapeutic application due to its variable properties including its antiviral effect.^11^ Star anise contains many compounds as flavonoids and polyphenols which have been investigated for its antiviral activity.^12^ This study aims to further explore the antiviral effect of combining *Lactiplantibacillus plantarum* and star anise against HSVs and VSV.

Consistent with Neli et. al which reported Pronounced virucidal activities of *L. plantarum*, we found that *L. plantarum* sonicate and supernatant could achieve 2.5 log reduction in HSV-1 titre.^13^ Another study reported significant reduction in virus yields in cutaneous HSV-1 infection model in mice.^14^

Our study showed that star anise extract could effectively reduce the titre of VSV and HSV. Fatma et. al reported that anise oil could inhibit the growth of bovine HSV-1 in cultured cells and induce 3 log reduction in virus titre. The combination of *L. plantarum* and star anise in our study has no superior effect over individual treatment with star anise or *L. plantarum*.

MX proteins, present in most vertebrates and coded by *MX* genes, have well known antiviral effects.^15^ Previous studies reported that MX proteins interfere with VSV replication either by inhibiting polymerase or affecting mRNA stability.^16^ MX proteins, induced by interferon, had been described as pan-herpesvirus restriction factor.^17^ Consistent with the result of viral replication assay, assessment of *MX* genes expression showed that treatment with star anise caused the greatest increase in *MX* genes in VSV infection while treatment with *L. plantarum* sonicate or supernatant caused the greatest increase in *MX* genes in HSV-1 infection. Wang et. al had previously described upregulated expression of interferon-stimulated genes in transmissible gastroenteritis virus infected mice treated with strain of *Lactobacillus plantarum*.^18^

## Conclusion

We demonstrated that star anise and *L. plantarum* have potent antiviral activities and they could be promising antiviral agents. Their inhibitory effect could be attributed to in-duction of *MX* genes expressions. Further studies are needed to explore underlying mechanism

## Author Contributions

All authors contributed to this work

## Institutional Review Board Statement

This project has received ethical approval from the Research Ethics Committee, Faculty of Science, South Valley University, Qena, Egypt, (REC-FScSVU) with ethics reference number (005/05/23).

## Data Availability Statement

The authors confirm that the data supporting the findings of this study are available within the article.

